# Genomic characterization of a pandrug-resistant *Klebsiella pneumoniae* belonging to the high-risk ST11 in the Brazilian Amazon region

**DOI:** 10.1101/2023.04.25.538267

**Authors:** Érica L. Fonseca, Sérgio M. Morgado, Fernanda S. Freitas, Nathalia S. Bighi, Rosângela Cipriano, Ana Carolina P. Vicente

**Affiliations:** Laboratório de Genética Molecular de Microrganismos, Instituto Oswaldo Cruz, FIOCRUZ, Rio de Janeiro, Brazil; São Domingos Hospital, São Luís do Maranhão, Brazil

**Author notes:** Corresponding author: Érica L. Fonseca.

**Keywords:** PDR, tigecycline resistance, colistin resistance, *ompK*, *acrAB*, untreatable bacteria

## Abstract

Pandrug-resistant (PDR) *K. pneumoniae* has been reported sporadically in many countries and remains rare in Brazil. The lack of genomic studies limits the comprehension of the determinants mostly involved with the PDR emergence in *K. pneumoniae*. This study aimed to unravel the main genetic determinants involved with the PDR background of a clinical ST11 *K. pneumoniae* recovered in the Brazilian Amazon region. The carbapenem-resistant Kp196 was submitted to WGS and its intrinsic and acquired resistome was assessed by CARD and comparison with wild-type genes. Kp196 resistome was composed of acquired resistance determinants and mutations in chromosomal genes. Among the formers, *bla*_CTX-M-15_ and *bla*_NDM-1_, *bla*_OXA-9_, *bla*_OXA-1,_ *aadA1, aacA4, strAB, aph(3’)-VI, aac(3)-IId, qnrS1, qnrB1, oqxAB, dfrA14, sul2, catB3* were found in the vicinity of mobile genetic elements, which could contribute to their spread. Kp196 colistin resistance was multifactorial and attributed to modifications in ArnT (M114L/V117I/R372K), PhoQ (D150G), and the *mgrB* disruption by IS*Kpn25*. Besides the presence of *qnr* and *oqxAB* genes, Kp196 also presented altered GyrA (S83I) and ParC (S80I). An *in-block* deletion in the repressor RamR, contributing to *acrAB* overexpression, and the presence of an enhanced-function AcrB variant (S966A), probably led to the Kp196 multidrug and tigecycline resistance. Insertions, *in-block* deletion, and missense mutations were involved with *ompK35-36-37* inactivation, also accounting for the Kp196 multidrug resistance, including carbapenems. The Kp196 PDR profile, especially the carbapenem resistance, was due to the accumulation of different mechanisms, in which modifications in housekeeping genes accounted for a more stable resistome.

## 1. Introduction

Pandrug resistance is related to the non-susceptibility to all agents in all antimicrobial categories considered approved and useful for treating an infection caused by a specific organism [1]. *Klebsiella pneumoniae* is featured by a remarkable propension for multidrug resistance acquisition, and infections caused by multidrug-(MDR) and extensively drug-resistant (XDR) strains are highly prevalent worldwide, while pandrug resistance (PDR) remains rare [2]. These MDR/XDR lineages are frequently carbapenem-resistant, and in this case, tigecycline and colistin remain the unique effective therapeutic choices [3]. Therefore, tigecycline and colistin co-resistance in carbapenem-resistant *K. pneumoniae* may result in apparently untreatable organisms, leading to a worrisome impact on clinical outcomes. Eventually, strains of the international high-risk clonal complex CC258 (ST11, ST437, and ST258) have presented the PDR profile. In Brazil, so far, PDR *K. pneumoniae* has only been reported in a few CC258 strains in the South/Southeast Brazilian regions [4-6], however, the genomic features involved with the PDR manifestation were rarely assessed. Here, the complete genome sequence of a clinical PDR *K. pneumoniae* strain, KP196, recovered in the Brazilian Amazon region was unraveled, and the main genetic determinants involved with the PDR background were revealed.

## 2. Material and Methods

The Kp196 was recovered in 2022 in a clinical setting in the Amazon Region (Maranhão). The antimicrobial susceptibility test (AST) was determined for all antibiotics considered for *Enterobacteriaceae* resistance classification [1], and interpreted according to the Clinical and Laboratory Standards Institute (CLSI) [7], and European Committee on Antimicrobial Susceptibility Testing (EUCAST) (for tigecycline and polymyxins) guidelines [8].

The Kp196 genome was obtained on the Illumina Hiseq 2500 using Nextera paired-end library kit for library construction. SPAdes assembler v3.15.2 was used for genome assembling [9]. Gene prediction/annotation was conducted with Prokka v1.14.6 [10]. Core genome MLST (cgMLST) was determined in Bacterial Isolate Genome Sequence Database (BIGSdb; http://bigsdb.pasteur.fr/klebsiella/). The Comprehensive Antibiotic Resistance Database (CARD) was used for antimicrobial resistance gene (ARG) prediction [11]. Plasmid replicon identification was conducted with the PlasmidFinder [12]. The deduced protein of each Kp196 chromosomal gene involved with resistance was compared with that of the wild-type reference strains *K. pneumoniae* NTUH-K2044 (**<u>NC_012731</u>.<u>1</u>**) and MGH 78578 (**<u>CP000647</u>**). The Kp196 genome sequence was deposited in the GenBank under accession no. **<u>JAQOSS000000000</u>** and with BioProject no. **<u>PRJNA926954</u>**.

## 3. Results and Discussion

The in vitro analyses revealed that Kp196 corresponded to a PDR strain (Table 1), and the cgMLST assigned it to the ST11 pandemic lineage. In spite of the high prevalence of this lineage in Brazil [13], this is the first report of a PDR ST11 in the country. In fact, PDR *K. pneumoniae* remains rare in Brazil, having only been reported in ST437 and ST258 restricted to the South/Southeast regions [4-6].

**Table 1.** Kp126 PDR phenotype

|  | MIC (mg/L) |  |  |  |  |  |  |  |  | Resistance profile determined by Disc-Diffusion<sup>c</sup> |
| --- | --- | --- | --- | --- | --- | --- | --- | --- | --- | --- |
|  | IPM<sup>a</sup> | MEM<sup>a</sup> | ETP<sup>a</sup> | DOR<sup>a</sup> | CZA<sup>a</sup> | C/T<sup>a</sup> | TGC<sup>a</sup> | CST<sup>b</sup> | PMB<sup>b</sup> |  |
| Kp196 | >32 | >32 | >32 | >32 | >256 | >256 | 0.75 | 8 | 4 | GEN, TOB, AMK, NET, CPT, TIM, TZP, CFZ, CXM, CTX, CAZ, FEP, FOX, CTT, CIP, SXT, ATM, AMP, AMC, SAM, CHL, FOF, TET, DOX, MIN |

<sup>a</sup>, MIC determined by E-Test method [7]. The new tigecycline breakpoints for resistance ( >0.5 mg/L) recently revised by EUCAST were applied [8].
<sup>b</sup>, MIC determined by broth microdilution method. Colistin and polymyxin B MIC breakpoints for resistance >2 mg/L [8].
<sup>c</sup>, AST determined by disk-diffusion method [7].
IPM, imipenem; MEM, meropenem; ETP, ertapenem; DOR, doripenem; CZA, ceftazidime/avibactam; C/T, ceftolozane/tazobactam; TGC, tigecycline; CST, colistin; PMB, polymyxin B; GEN, gentamicin; TOB, tobramycin; AMK, amikacin; NET, netilmicin; CPT, ceftaroline; TIM, ticarcillin/clavulanic acid; TZP, piperacillin/tazobactam; CFZ, cefazolin; CXM, cefuroxime; CTX, cefotaxime; CAZ, ceftazidime; FEP, cefepime; FOX, cefoxitin; CTT, cefotetan; CIP, ciprofloxacin; SXT, trimethoprim/sulfamethoxazole; ATM, aztreonam; AMP, ampicillin; AMC, amoxicillin/clavulanic acid; SAM, ampicillin/sulbactam; CHL, chloramphenicol; FOF, fosfomycin; TET, tetracycline; DOX, Doxycycline; MIN, minocycline.

The PDR phenotype was in accordance with the Kp196 expressive resistome, which was composed of genes associated with resistance to β-lactams (*bla*_SHV-11_, *bla*_CTX-M-15_, *bla*_OXA-9_, *bla*_OXA-1,_ bla_TEM-1_), aminoglycosides (*aadA1, aacA4, strAB, aph(3’)-VI, aac(3)-IId*), carbapenems (*bla*_NDM-1_) fluoroquinolones (*qnrS1, qnrB1, oqxAB*), trimethoprim (*dfrA14*), sulfonamides (*sul2*), tetracycline (*tetD*), fosfomycin (*fosA5*) and chloramphenicol (*catB3*). All these genes were flanked or in the vicinity of insertion sequences and plasmid-related genes. In fact, Kp196 harboured *repA, repB* and *repE* genes from IncFIB and IncR plasmids. The exception was the *tetD, fosA5*, and *bla*_SHV-11_, which were chromosomally encoded.

Regarding the intrinsic mechanisms, mutations were observed in genes involved with resistance to fluoroquinolones (*gyrA, parC*), colistin (*mgrB, arnT, phoQ*), tigecycline (*ramR*), and multiple drugs including carbapenems and cephalosporins (*acrB, ompK35, ompK36*, and *ompK37*). Ciprofloxacin is effective and widely used for treating ESBL-producing *K. pneumoniae* infections. The Kp196 presented substitutions in the quinolone resistance-determining region (QRDR) of GyrA (S83I) and ParC (S80I), which are involved with ciprofloxacin resistance emergence in *K. pneumoniae*.

Colistin resistance in *K. pneumoniae* is mainly associated with modifications in *pmrAB, phoPQ, mgrB*, and *arnT* genes [14]. Among these genes, substitutions were found in the deduced protein of ArnT (M114L, V117I, and R372K), involved with *pmrA* transcription, and in PhoQ (D150G). Besides, the *mgrB* was disrupted by IS*Kpn25* at the nucleotide position 133, leading to the production of a truncated and inactive MgrB protein, probably contributing to Kp196 colistin resistance due to *phoPQ* derepression [14]. Interestingly, this same alteration was previously found in colistin-resistant ST258 *K. pneumoniae* from Greece and Brazil [15], indicating that this region might be a hotspot for IS*Kpn25* insertion. This IS additionally carried *bla*_TEM-1_, *aac(3)-IId*, and a complete Restriction modification System (RMS), also contributing to β-lactams and aminoglycosides resistance, and to host protection from foreign DNA infection. Therefore, the Kp196 colistin resistance could be associated with the accumulation of multiple alterations in chromosomal genes (*mgrB, arnT*, and *phoQ*). In this case, even upon restoration of the canonical function by reversal mutations in one of these genes, Kp196 would retain the colistin resistance (Table 2).

**Table 2.**
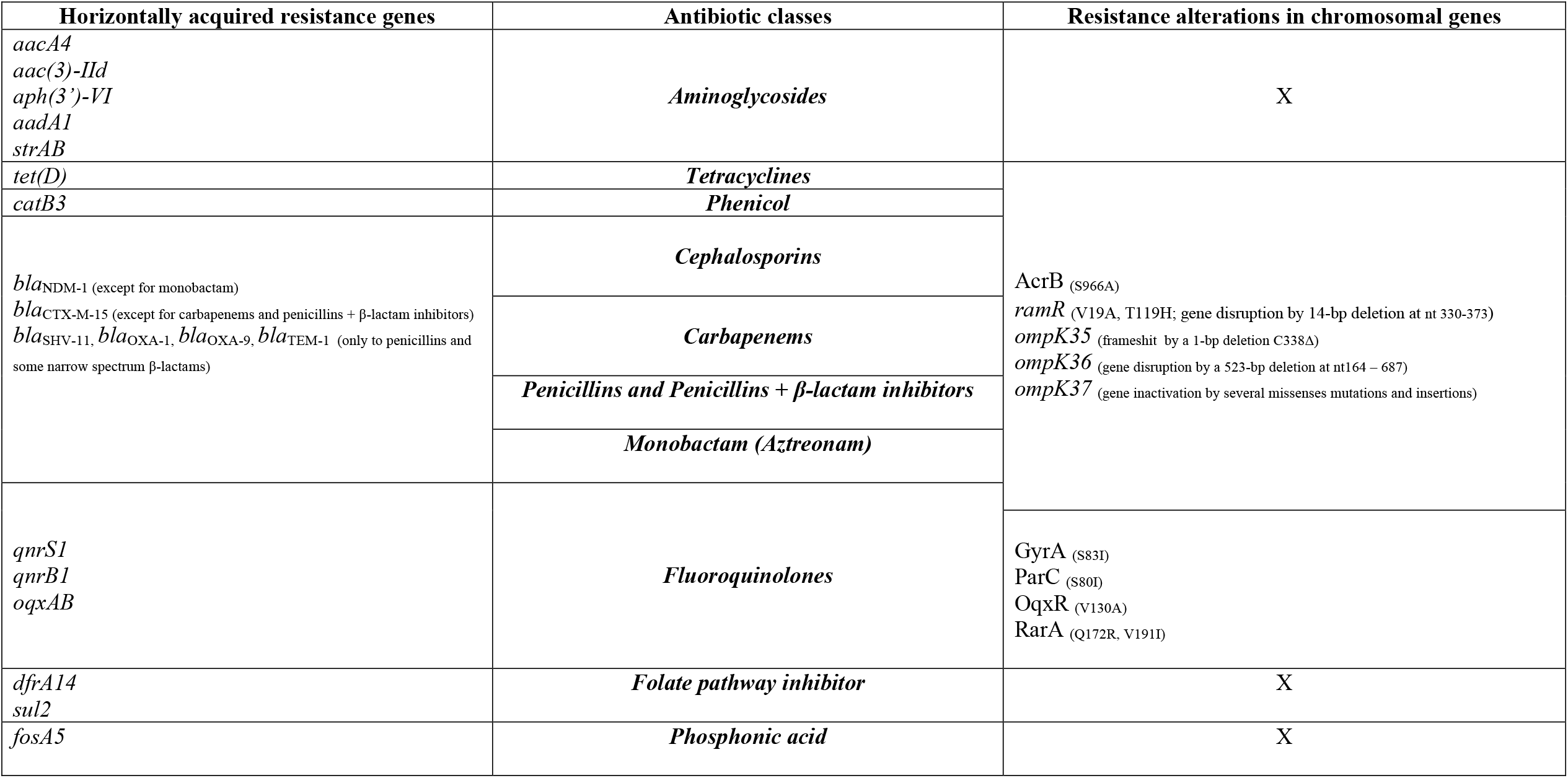
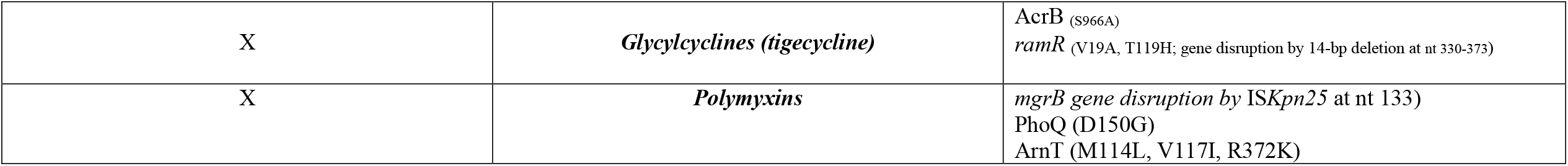
Acquired and intrinsic resistance mechanisms involved with the Kp196 PDR phenotype

In spite of several tigecycline resistance mechanisms already described, *K. pneumoniae* tigecycline-resistant strains remain rare [15]. Among these mechanisms, efflux pump overexpression (*acrAB* and *oqxAB*) due to alterations in their regulatory genes (*ramR, ramA, soxR, soxS, marA, marR, acrR, oqxR, rarA*) is the most common [16]. From all the aforementioned regulatory genes, only *ramR* (*ramA* repressor), *oqxR* and *rarA* (*oqxAB* repressor and activator, respectively) were altered in Kp196. The RamR presented two amino acid modifications (V19A and T119H) and a 14 bp-deletion downstream the nt 330 was present in this gene, leading to a frameshift. This *in-block* deletion probably generated an inactivated RamR, resulting in *ramA* derepression and, consequently, to *acrAB* overexpression. The substitutions found in RarA (Q172R and V191I) have not been described yet, while the OqxR presented the V130A alteration that had already been found in tigecycline-susceptible strains [16]. Therefore, the *ramR* alterations were probably the main tigecycline and multidrug resistance determinant in Kp196 (Table 2).

Kp196 harboured the S966A AcrB variant, which is involved with the increment of drug transport efficiency, conferring an increased ability to persist/resist its substrate antibiotics when overexpressed [17]. Since *acrAB* is also involved with resistance to other tetracyclines, fluoroquinolones, erythromycin, β-lactams, chloramphenicol, and also carbapenems [18-20], the *acrAB* overexpression with an enhanced-function AcrB variant may also contribute with the remarkable Kp196 multidrug resistance phenotype.

In *K. pneumoniae*, loss of the two major outer membrane porins OmpK35 and OmpK36 enhances the multidrug resistance in ESBL-producing strains, increasing resistance to carbapenems, broad-spectrum cephalosporins, fluoroquinolones, tetracycline, and chloramphenicol [21]. In Kp196, the *ompK35* suffered a deletion at nucleotide 338 resulting in a frameshift, while an *in-block* deletion from nucleotide 164 to 687 disrupted *ompK36*. The *ompK37* is normally expressed only in *ompK35-36*-deficient strains, slightly influencing carbapenem resistance [21]. However, in addition to *ompK35*/*36*, the *ompK37* of Kp196 was also altered, presenting a set of SNPs and insertions along the gene, leading to a defective porin. Therefore, all three *K. pneumoniae* major porins were inactivated in Kp196, contributing significantly to multidrug resistance in this strain. Finally, considering the clinical relevance of carbapenem resistance, this study stressed the multifactorial and overrepresented mechanisms in Kp196, which comprised the presence of *bla*_NDM-1_ and alterations of several intrinsic genes, such as *acrAB, ompK35-36-37*.

Interestingly, the unique genomic studies on CC258 *K. pneumoniae* PDR strains in Brazil demonstrated a different resistome composition compared to Kp196, considering both the intrinsic and acquired resistance determinants involved with PDR manifestation [4,6]. Besides, in both studies, the PDR phenotype was mainly due to the presence of acquired resistance genes.

## 4. Conclusion

Here, a clinical strain was described in the Amazon region likely presenting a more stable resistome, since multiple mutations in chromosomal genes, which are not easily lost as the acquired resistance determinants, importantly contributed to the observed PDR phenotype.

## Ethical Approval

Not required.

